# Re-Evaluating One-step Generation of Mice Carrying Conditional Alleles by CRISPR-Cas9-Mediated Genome Editing Technology

**DOI:** 10.1101/393231

**Authors:** Channabasavaiah Gurumurthy, Rolen Quadros, John Adams, Pilar Alcaide, Shinya Ayabe, Johnathan Ballard, Surinder K. Batra, Marie-Claude Beauchamp, Kathleen A Becker, Guillaume Bernas, David Brough, Francisco Carrillo-Salinas, Ruby Dawson, Victoria DeMambro, Jinke D’Hont, Katharine Dibb, James D. Eudy, Lin Gan, Jing Gao, Amy Gonzales, Anyonya Guntur, Huiping Guo, Donald W. Harms, Anne Harrington, Kathryn E. Hentges, Neil Humphreys, Shiho Imai, Hideshi Ishii, Mizuho Iwama, Eric Jonasch, Michelle Karolak, Bernard Keavney, Nay-Chi Khin, Masamitsu Konno, Yuko Kotani, Yayoi Kunihiro, Imayavaramban Lakshmanan, Catherine Larochelle, Catherine B. Lawrence, Lin Li, Volkhard Lindner, Xian-De Liu, Gloria Lopez-Castejon, Andrew Loudon, Jenna Lowe, Loydie Jerome-Majeweska, Taiji Matsusaka, Hiromi Miura, Yoshiki Miyasaka, Benjamin Morpurgo, Katherine Motyl, Yo-ichi Nabeshima, Koji Nakade, Toshiaki Nakashiba, Kenichi Nakashima, Yuichi Obata, Sanae Ogiwara, Mariette Ouellet, Leif Oxburgh, Sandra Piltz, Ilka Pinz, Moorthy P. Ponnusamy, David Ray, Ronald J. Redder, Clifford J Rosen, Nikki Ross, Mark T. Ruhe, Larisa Ryzhova, Ane M. Salvador, Radislav Sedlacek, Karan Sharma, Chad Smith, Katrien Staes, Lora Starrs, Fumihiro Sugiyama, Satoru Takahashi, Tomohiro Tanaka, Andrew Trafford, Yoshihiro Uno, Leen Vanhoutte, Frederique Vanrockeghem, Brandon J. Willis, Christian S. Wright, Yuko Yamauchi, Xin Yi, Kazuto Yoshimi, Xuesong Zhang, Yu Zhang, Masato Ohtsuka, Satyabrata Das, Daniel J. Garry, Tino Hochepied, Paul Thomas, Jan Parker-Thornburg, Antony D Adamson, Atsushi Yoshiki, Jean-Francois Schmouth, Andrei Golovko, William R. Thompson, KC. Kent Lloyd, Joshua A. Wood, Mitra Cowan, Tomoji Mashimo, Seiya Mizuno, Hao Zhu, Petr Kasparek, Lucy Liaw, Joseph M. Miano, Gaetan Burgio

**Affiliations:** Mouse Genome Engineering Core Facility, Vice Chancellor for Research Office, University of Nebraska Medical Center, Omaha, NE, USA; Developmental Neuroscience, Munroe Meyer Institute for Genetics and Rehabilitation, University of Nebraska Medical Center, Omaha, NE, USA; Texas A&M Institute for Genomic Medicine (TIGM), Texas A&M University, College Station, TX 77843, USA; Department of Immunology, Tufts University School of Medicine, Boston, USA; RIKEN BioResource Research Center, Tsukuba, Ibaraki 305-0074, Japan; Department of Biochemistry and Molecular Biology, University of Nebraska Medical Center, Omaha, NE, USA; Departments of Anatomy and Cell Biology, Human Genetics and Pediatrics, Research Institute McGill University Health Center (RI-MUHC), Montreal, Canada; Maine Medical Center Research Institute (MMCRI),Scarborough, ME, USA; Transgenesis and Animal Modeling Core Facility, Centre de Recherche du Centre Hospitalier Universitaire de Montreal (CRCHUM), Montreal, Canada; Division of Neuroscience and Experimental Psychology, School of Biological Sciences, Faculty of Biology, Medicine and Health, Manchester Academic Health Science Centre, University of Manchester, AV Hill Building, Oxford Road, Manchester, M13 9PT, U.K; South Australian Health & Medical Research Institute and Department of Medicine, University of Adelaide, Australia; Transgenic mouse core facility, VIB Center for Inflammation Research, Ghent, Belgium; Department of Biomedical Molecular Biology, Ghent University, Ghent, Belgium; Unit of Cardiac Physiology, School of Medical Sciences, Manchester Academic Health Science Center, University of Manchester, UK; High-Throughput DNA Sequencing and Genotyping Core Facility, Vice Chancellor for Research Office, University of Nebraska Medical Center, Omaha, USA; University of Rochester Medical Center, Rochester, NY 14642, USA; Division of Evolution and Genomic Sciences, School of Biological Sciences, Faculty of Biology, Medicine and Health, Manchester Academic Health Science Centre, University of Manchester, UK; Transgenic Unit core facility, Faculty of Biology, Medicine and Health, University of Manchester, Manchester, UK; Department of Basic Medicine, Division of Basic Medical Science and Molecular Medicine, School of Medicine, Tokai University, 143, Shimokasuya, Isehara, Kanagawa 259-1193, Japan; Department of Medical Data Science, Osaka University Graduate School of Medicine, Japan; The University of Texas, MD Anderson Cancer Center, Houston, TX, USA; Division of Cardiovascular Sciences, School of Medical Sciences, Faculty of Biology, Medicine and Health, The University of Manchester AND Manchester Heart Centre, Manchester University NHS Foundation Trust, Manchester Academic Health Science Centre, Manchester, UK; Department of Frontier Science for Cancer and Chemotherapy, Osaka University Graduate School of Medicine, Osaka, Japan; The Institute of Experimental Animal Sciences, Osaka University Graduate School of Medicine, Osaka, Japan; Centre de Recherche du Centre Hospitalier Universitaire de Montreal (CRCHUM), Montreal, Canada; Children’s Research Institute Mouse Genome Engineering Core, University of Texas Southwestern Medical Center, Dallas, TX 75390, USA; Manchester Collaborative Centre for Inflammation Research (MCCIR), School of Biological Sciences, Faculty of Biology, Medicine and Health, The University of Manchester, Manchester, UK; Centre for Biological Timing, School of Medical Sciences, Faculty of Biology, Medicine and Health, University of Manchester, Manchester, UK; Center for Matrix Biology and Medicine, Graduate School of Medicine, Tokai University, Isehara, Kanagawa, 259-1193, Japan; Department of Molecular Life Science, Division of Basic Medical Science and Molecular Medicine, School of Medicine, Tokai University, 143, Shimokasuya, Isehara, Kanagawa 259-1193, Japan; Laboratory of Molecular Life Science, Foundation for Biomedical Research and Innovation, at Kobe, Japan; Department of Laboratory Animal Science, Support Center for Medical Research and Education, Tokai University, 143, Shimokasuya, Isehara, Kanagawa 259-1193, Japan; Oxford Centre for Diabetes, Endocrinology and Metabolism, University of Oxford, Oxford, OX37LE, UK; Mouse Biology Program, University of California, Davis, USA; Laboratory of Transgenic Models of Diseases and Czech Centre for Phenogenomics, Institute of Molecular Genetics of the Czech Academy of Sciences, Czech Republic; College of Osteopathic Medicine, Marian University, Indianapolis, IN 46222, USA; Laboratory Animal Resource Center, University of Tsukuba, Japan; Department of Gastroenterology and Metabolism, Nagoya City University Graduate School of Medical Sciences, Nagoya, Japan.; School of Health and Human Sciences, Department of Physical Therapy, Indiana University, Indianapolis, IN 46202, USA; Lillehei Heart Institute Regenerative Medicine and Sciences Program, University of Minnesota, Minneapolis, MN, USA; Paul and Sheila Wellstone Muscular Dystrophy Center, University of Minnesota, Minneapolis, MN, USA; Dept. of Surgery, School of Medicine, University of California, Davis, Davis, USA; McGill Integrated Core for Animal Modeling (MICAM), Montreal, Canada; Department of Immunology and Infectious Disease, the John Curtin School of Medical Research, the Australian National University, Canberra, Australia

## Abstract

CRISPR-Cas9 gene editing technology has considerably facilitated the generation of mouse knockout alleles, relieving many of the cumbersome and time-consuming steps of traditional mouse embryonic stem cell technology. However, the generation of conditional knockout alleles remains an important challenge. An earlier study reported up to 16% efficiency in generating conditional knockout alleles in mice using 2 single guide RNAs (sgRNA) and 2 single-stranded oligonucleotides (ssODN) (2sgRNA-2ssODN). We re-evaluated this method from a large data set generated from a consortium consisting of 17 transgenic core facilities or laboratories or programs across the world. The dataset constituted 17,887 microinjected or electroporated zygotes and 1,718 live born mice, of which only 15 (0.87%) mice harbored 2 correct *LoxP* insertions in *cis* configuration indicating a very low efficiency of the method. To determine the factors required to successfully generate conditional alleles using the 2sgRNA-2ssODN approach, we performed a generalized linear regression model. We show that factors such as the concentration of the sgRNA, Cas9 protein or the distance between the placement of *LoxP* insertions were not predictive for the success of this technique. The major predictor affecting the method’s success was the probability of simultaneously inserting intact proximal and distal *LoxP* sequences, without the loss of the DNA segment between the two sgRNA cleavage sites. Our analysis of a large data set indicates that the 2sgRNA–2ssODN method generates a large number of undesired alleles (>99%), and a very small number of desired alleles (<1%) requiring, on average 1,192 zygotes.

## Introduction

Many inherited diseases are caused by defective genes. A better understanding of the mechanisms of these defects is critical to obtaining precise diagnoses and finding new therapeutics. Gene inactivation through knockout alleles in model organisms such as flies, worms, zebrafish and mice provides invaluable insights into mechanisms of gene function and disease [1]. However important challenges remain to successfully analyze the phenotypic impact of knockout genes in adult model organisms as over 30% of the genes in mice are essential for development and cause embryo lethality or neonatal subviability when deleted [2]. To overcome lethal phenotypes in gene-knockout models, conditional knockout (cKO) strategies have emerged [3]. cKO models usually involve insertion of *LoxP* sites in introns flanking critical exon/s or (less commonly) in intergenic regions or flanking regulatory regions such as promoters and enhancers. When crossed with a *Cre* recombinase expressing driver mouse, the *Cre* enzyme recognizes *LoxP* sequences and removes the intervening sequence. This leads to functional inactivation of the targeted gene in only the cells where the *Cre* is expressed and capable of targeting the DNA [3]. Generating a cKO mouse previously required the use of embryonic stem (ES) cell-based homologous recombination in combination with embryo manipulation, microinjection (MI), and assisted reproduction technologies (ART) [4]. These techniques were established in the 1980s and are still being used as gold-standard methods. Based on this technology, large-scale efforts such as the KnockOut Mouse Project (KOMP) [5] and the European Conditional Mouse Mutagenesis (EUCOMM) Program [6] have designed thousands of gene targeted constructs in ES cells for over 90% of coding genes. Using the ES cell clones, about 25% of mouse genes have been converted into cKO mice, all readily available and accessible in public repositories [7].

The recent emergence of genome editing technologies such as ZFN, TALENS and CRISPR-Cas9 enables an improvement in efficiency of gene targeting and has considerably facilitated the generation of genetically-engineered animal models based on homology directed repair of donor constructs in mouse zygotes [8]. Endonucleases, particularly Class 2 CRISPR systems, generate a precise double strand break (DSB) in the DNA under a chimeric single guide RNA (sgRNA) [9]. The DSB leads to error-prone, non-homologous end joining (NHEJ) repair or the precise homology-directed repair (HDR) under the guidance of a repair template [8]. In an earlier study, a high success rate (16%) of targeting *LoxP* sites in *cis* was reported by using 2 sgRNAs and 2 single-stranded oligonucleotides (ssODN) containing *LoxP* sites (2sgRNA-2ssODN) flanking a targeted critical exon (Figure 1) [10].

**Figure 1:**
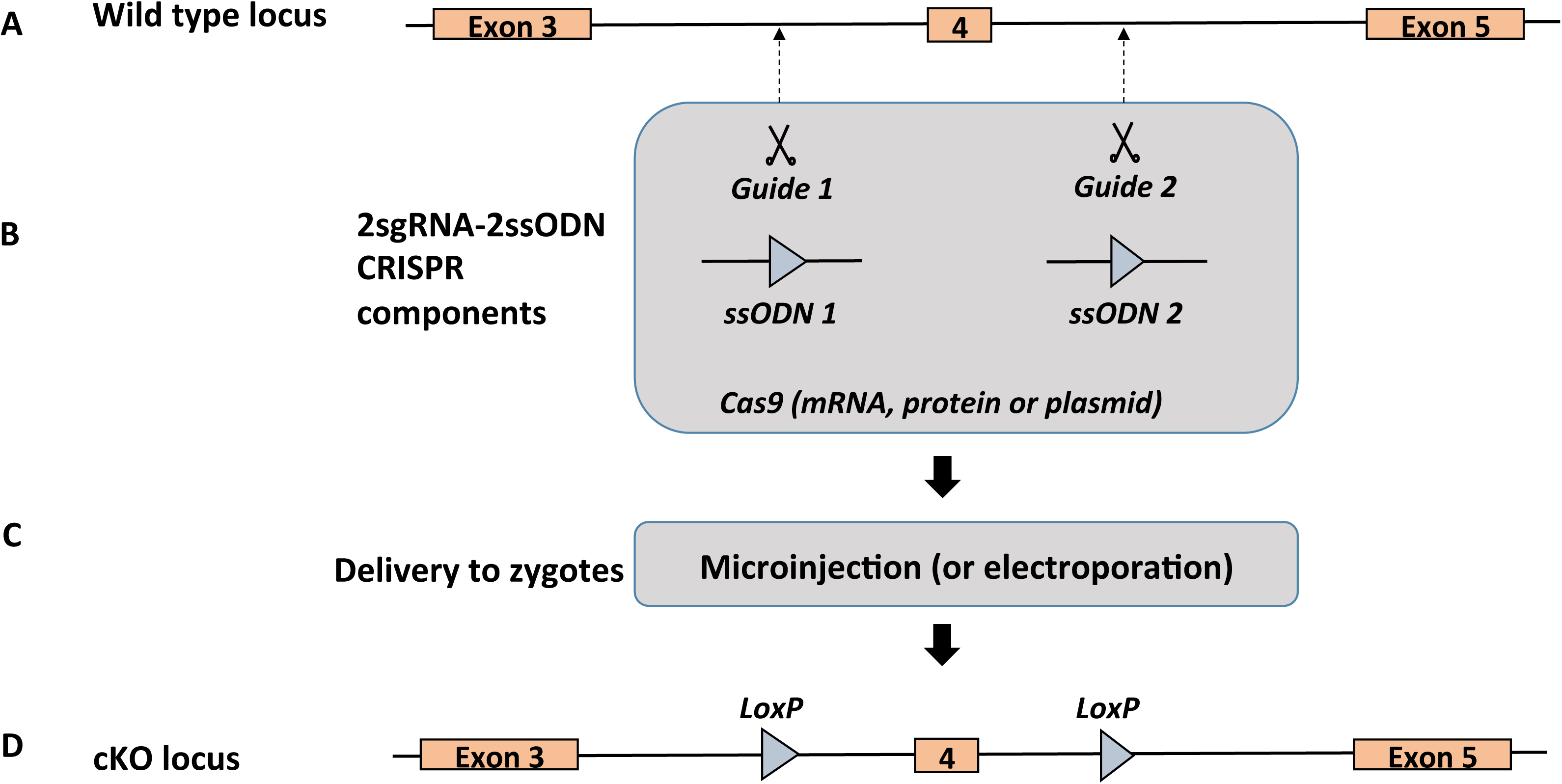
Schematic of 2sgRNA-2ssODN CRISPR method of creating conditional knockout alleles. (A) Wild type locus showing exons 3, 4 and 5 of a hypothetical gene where exon 4 is chosen as a target exon for inserting *LoxP* sites. The guide 1 and the guide 2 target introns 3 and 4 respectively. (B) CRISPR components showing 2sgRNA-2ssODN donors and a Cas9 source. (C) Delivery of CRISPR components into one-cell stage zygotes via microinjection (n=17,867) or electroporation (n=330). (D) The conditional knockout (cKO) allele showing target exon (#4) with flanking *LoxP* sites.

We sought to investigate the efficiency of the 2sgRNA-2ssODN method for the generation of cKO alleles. We describe here for the first time a global community effort from a consortium of over a dozen laboratories, transgenic core facilities and programs across the world to evaluate the efficiency of generating cKO alleles using the 2sgRNA-2ssODN approach. We surveyed over 50 loci and over 17,000 microinjected or electroporated zygotes using this method, which enabled robust statistical power to evaluate the efficiency of the technique. In contrast to the earlier report [10], we find this method does not efficiently produce cKO alleles. Rather, it generally results in a series of undesired editing events at the cleavage sites which occur nearly 100 fold higher rate than the precise insertion, in *cis*, of the two *LoxP* sites.

## Material and methods

### Ethical statement

All experiments were approved from the respective Institutional Animal Care and Use Committees in the USA and Ethics Committees in Australia, Belgium, Czech Republic, Japan, Spain and UK according to guidelines or code of practice from the National Institute of Health in the USA, the National Health and Medical Research Council (NHMRC) in Australia, Animals (Scientific Procedures) Act 1986 in UK or MEXT (Ministry of Education, culture, sports, Science and Technology), MHLW (The Ministry of Health, Labor and Welfare) in Japan, the central commission for Animal Welfare (CCAW) in Czech Republic, the Canadian Council on Animal Care (CCAC) in Canada, the National Ethics Code from the Royal Belgian (Flemish) Academy of Medicine in Belgium, and the European code of Conduct for Research Integrity from All European Academies.

### *Mecp2* gene targeting using CRISPR-Cas9

*Mecp2* left single chimeric guide RNAs (sgRNA) 5’-CCCAAGGATACAGTATCCTA-3’ and *Mecp2* right sgRNA 5’-AGGAGTGAGGTCTAGTACTT-3’ target sites were designed as described in Yang et al [10]. Ultramer Oligonucleotides (Integrated DNA Technologies, Coralville, IA) were designed with sequences to T7 promoter for *in vitro* transcription, DNA target region, and chimeric RNA sequence. Complimentary oligos for each target sequence were annealed at 95°C for 5 mins and the temperature was reduced 0.20°C/second to 16°C using a PCR machine (BioRad T100) before use as template for sgRNA synthesis. sgRNAs were synthesized with the HiScribe™ T7 Quick High Yield RNA Synthesis Kit (New England Biolabs). Cas9 mRNA was obtained from Life Technologies or in-vitro transcribed from a Chimeric pX330-U6-Chinmeric-BB-CBh-hSpCas9 expression plasmid obtained from Addgene repository (Plasmid 42230; donation from Zhang laboratory).

### SgRNA design

SgRNAs were designed using available online tools such as CRISPOR, Chop-Chop or CCTop [11, 12]. SgRNAs were cloned into pX330 and *in vitro* transcribed[13-15], or synthesized and annealed [16]. Cas9 mRNA or protein was purchased, *in vitro* transcribed or purified in house. Cas9 protein was complexed with thesgRNA or crRNA and the tracrRNA [17] and then mixed with the ssODN prior to microinjection. Concentrations and site of injection for Cas9 protein or mRNA, sgRNA, and template repairs for each locus are indicated in Supplementary Table 1.

### Mouse husbandry and zygote microinjection and electroporation

Mice were purchased from various sources and maintained under specific pathogen-free conditions. Mice were maintained under 12/12 hr light cycle and food and water were provided *ad libitum*. Three to five week-old females were superovulated by intraperitoneal injection of Pregnant Mare Serum Gonadotropin (5IU) followed by intraperitoneal injection of Human Chorionic Gonadotropin hormone (5IU) 48 hours later. Superovulated females were mated with 8 to 20 week-old stud males. The mated females were euthanized the following day and the zygotes were collected from their oviducts. Cytoplasmic or pronuclear injections were performed under an inverted microscope, associated micromanipulators, and a microinjection apparatus. Electroporation of the embryos were performed with an electroporation device using a cuvette or 1mm plate electrodes with the following parameters: 30 V square wave pulses with 100 ms interval using a BioRad electroporator device or 4 poring pulses (40 V, 3.5 ms, interval 50 ms, 10% voltage decay + polarity) followed by 5 transfer pulses (5 V, 50!ms, interval 50 ms, 40% voltage decay, alternating + and - polarity) using a NEPA21 electroporator device. Microinjected or electroporated zygotes were either surgically transferred into the ampulla of pseudo-pregnant females or cultured overnight at 37°C and then surgically transferred at the 2-cell stage of development.

### Genotyping

DNA extraction was performed on ear punch or tail tip from mouse pups over 15 days using a DNA extraction kit according to the manufacturer instructions. Primers were designed to amplify the regions encompassing the integrated *LoxP* sequence. PCR was performed using Taq polymerase under standard PCR conditions. The PCR products were then purified with ExoSAP-IT1 or a PCR Clean-Up System kit according to the manufacturer’s instructions. Sanger sequencing was performed in core facilities. To identify *LoxP* insertions, as a general practice at all centers, the two target sites were amplified individually to look for increase in the amplicon size, which occurs if *LoxP* sites are inserted successfully. If the *LoxP* insertion was not observed in this first set of PCR analyses, the samples were declared negative, and in many such cases the samples were not analyzed further (as the end goal of the project, ie., generation of floxed allele was not met). In some cases, such samples were also sequenced to assess *indels* to understand if the guides were successful in cleaving the target site. In some cases, the entire regions encompassing both the guide cleavage sites were amplified to assess for deletions between the cleavage sites.

### Statistics

To determine the statistical differences between proportions or means, we performed a Fisher Exact test or a Kruskal Wallis sum rank test. A Generalized linear model calculation was performed with success of the 2sgRNA-2ssODN method as a response. Predictive variables were: efficiency of the sgRNA, probability of *LoxP* insertions in 5’ and 3 (5’_*LoxP* and 3’_*LoxP*), simultaneous insertion of the 2 *LoxP* sites (interaction between 5’_*Loxp* and 3’_*LoxP*) Cas9 mRNA, protein, plasmid and ssODN concentrations and distance between distal and proximal target sites. Variance for each predictor was determined from the diagonal of the variance-covariance matrix. Effect sizes and type II error were determined using Cohen effect size d statistics and power calculation. All statistical analyses were performed using Rstudio v1.1.423. Results were considered statistically significant at p<0.05.

## Results

### *Mecp2* gene targeting in blastocysts

To assess the efficiency of the technique and compare to previously published results [10], we reproduced an experiment on *Mecp2* gene, essential for DNA methylation during development using the same sgRNAs and ssODNs as previously described in the original report [10]. Three independent centers at the Australian National University in Australia (ANU), University of Nebraska Medical Center in the USA (UNMC) and the Czech Centre for Phenogenomics in Czech Republic (IMG) performed these experiments on C57BL/6N inbred strain of mice. We evaluated the success rate of the 2sgRNA-2ssODN method in blastocysts for *Mecp2* (Table 1). Using a concentration mix of 20 ng/µl of Cas9 mRNA, 20 ng/µl of in-vitro transcribed sgRNA, and 10 ng/µl of ssODN, we observed no successful targeting (i.e., correct insertion of 2 *LoxP* sites in *cis* configuration) even though both sgRNAs cleaved target DNA as indicated by the presence of *indels* or integration of a *LoxP* site at the desired location, which varied from 13% to 34% (Table 1).

**Table 1:** Summary of the edited blastocysts for *Mecp2* gene from three different centers.

|  | **Zygotes injected** | **Blastocysts genotyped** | **Correctly targeted** | **Incorrectly targeted at the 5' site (%)** | **Incorrectly targeted at the 3' site (%)** |
| --- | --- | --- | --- | --- | --- |
| Australian National University (ANU) Australia | 106 | 51 | 0 | 11 *indels* and 6 *LoxP* correctly inserted (33%) | 6 *indels* and 1 *LoxP* correctly inserted (13%) |
| University of Nebraska Medical Center (UNMC) USA | 80 | 70 | 0 | 14 *indels* and 1 *LoxP* correctly inserted (34%) | 21 *indels* (30%) |
| Czech Centre for Phenogenomics, Czech republic (BIOCEV/IMG) | 40 | 28 | 0 | 8 *indels* and 1 *LoxP* correctly inserted (32%) | 5 *indels* (18%) |

Interestingly we noted the occasional presence of mutations within *LoxP* sites indicating illegitimate repair events at the target site. The frequency of successful targeting of two *LoxP* sites *in cis* was previously reported to be 16% [10], which we failed to achieve. One possible explanation is the mouse genetic background influences the likelihood of ssODN integration. This variance could also be explained by an inherently low probability to successfully replace 2 genomic loci in *cis*, the lack of efficiency of the sgRNA, or the relatively low sample size.

### A global survey of the generation of conditional alleles using 2sgRNA-2ssODN method

To better understand how to successfully generate conditional alleles using the 2sgRNA– 2ssODN approach and to assess its efficiency, we evaluated this method on 56 additional genes and two intergenic regions of the mouse genome from a consortium of 17 institutions across Australia, Belgium, Japan, USA, UK, Czech Republic and Canada. A majority of attempts were performed on a C57BL/6J background (39) whereas 18 projects used C57BL/6N background and 3 additional ones used a hybrid mouse background (B6C3HF1, B6SJLF1, FVBCD1F1). We assessed whether the mouse background strain would have an impact over the success of the method using Fisher Exact test statistics. We failed to find such evidence in our data (Fisher exact test, p = 0.74). Out of the 56 targeted loci (49 microinjected and 7 electroporated), 21 were ranked as essential genes based on early embryonic or postnatal lethality of the homozygous knockout mice according to mouse genome database http://www.informatics.jax.org [18]. Different knockout mice from 18 out of 56 targeted loci were described viable to adulthood as homozygous mice and 17 loci were unknown. Together this indicates the repartition between putative essential and non-essential targeted gene was in equal frequency (Fisher exact test, p = 0.76). The distance between sgRNA varied from 250 bp to 1.1 Mb with a median of 2 Kb. Single exons to entire genes or regulatory genomic regions (Supplementary Table 1) were floxed. We investigated whether the distance between sgRNA is critical for the likelihood of success of the 2sgRNA-2ssODN method. We failed to find such evidence in our data set (Kruskal Wallis rank sum test, chi-squared = 32, p=0.42), although the sample size was too low to form a conclusion (Cohen’s effect size d = 0.40 with power 1-beta = 0.27). Among the microinjected zygotes in 53 Loci, significantly higher number of zygotes were microinjected in the pronucleus alone (26/53) than the cytoplasm alone (10/53) or pronucleus and cytoplasm (17/53) (Fischer exact test p = 0.004), which is consistent with the current practice in most mouse transgenic core facilities (Figure 2A). Various forms of CRISPR reagents (sgRNA, Cas9 and ssODN), were microinjected or electroporated to generate the models (Supplementary Table 1). Consistent with the general practice in mouse transgenic facilities from 2013 to 2016 using CRISPR-Cas9 gene editing technology, the majority of the reagents were delivered in 59 Loci (49 unique loci microinjected, 3 different designs for one loci and 7 electroporated Loci) in the form of in-vitro transcribed mRNA (35/59) at various concentrations varying from 10 ng/µl to 100 ng/µl of Cas9 mRNA (Figure 2B) and from 10 ng/µl to 50 ng/µl sgRNA. ssODN were delivered at a concentration varying from 10 ng/µl to 200 ng/µl. In 18 instances, Cas9 was delivered as protein with a concentration varying from 10 ng/µl to 75 ng/µl. Interestingly for 6 loci, Cas9 and sgRNAs were delivered in the form of a chimeric sgRNA-SpCas9 plasmid (pX330) at a concentration of 5 ng/µl. We sought to determine whether the forms of reagent delivery such as plasmid, ribonucleoprotein (RNP) or mRNA would have an effect on the overall efficiency in targeting using the 2sgRNA-2ssODN method. We failed to find such evidence (Fisher exact test p = 1). We therefore hypothesized that the success in generating floxed alleles using the 2sgRNA-2ssODN approach may depend on factors such as: (i) sgRNA efficiency, (ii) simultaneity in *LoxP* insertion, or; (iii) the concentration of the Cas9, sgRNA and ssODN reagents. To get insight on these possibilities, we further analyzed data from the 56 loci (Supplementary Table 2, 4 and 5). Note that the offspring for 54 loci were analyzed post-natal stage (Supplementary Tables 2 and 4) whereas 2 loci were analyzed at the blastocyst stage (Supplementary Table 4). Out of 17,887 (17,557 microinjected and 330 electroporated; see details below) zygotes, 12,764 (71.4%) zygotes were surgically transferred into recipient females. The recipient females gave birth to 1,718 pups (9.6% of the microinjected/electroporated zygotes). As a general practice, at all centers, the mice were first analyzed by PCR to observe the putative insertion of the *LoxP* sites at both the sites; the animals were declared negative if genotyping did not reveal the presence of the desired allele. In some cases the loci were further analyzed to assess guide cleaving activity. Of the 1,684 founder mice, 659 (39%) showed some type of editing (*indels* and/or substitutions), 235 (14%) and 144 (9%)mice harbored a single *LoxP* insertion or deletions between the two cleavage sites, respectively (Figure 2C). The mice for 25 (of the 56) loci were further assessed for additional events including large deletions (Figure 2C). Of the 487 founder mice analyzed (from those 25 loci), 41%, 11% and 2.7% samples contained *indels,* single *LoxP* insertions or large deletions respectively (Figure 2D). From the 1,684 animals analyzed, only 15 mice (0.87%) were correctly targeted with intact *LoxP* sites in the *cis* configuration (Supplementary Table 5). Out of 56 loci only 11 loci were successfully targeted (19.6%). The average number of zygotes needed to generate 1 correctly targeted animal was 1,192. The essentiality of the genes had no impact on the likelihood of success of the 2sgRNA-2ssODN technique (4/23 success in targeting for embryonic or postnatal lethality versus 5/18 for viable homozygous mice and 2/15 for unknown embryonic or postnatal lethality, Fisher exact test p = 0.27). We also noted from our data, among the 56 loci analyzed 14% loci showed deletions between two target sites for Cas9 cleavage. We also noted a relatively high occurrence of single *LoxP* insertions for > 20% of the mice genotyped (from all the loci) and few instances of *trans LoxP* insertions (on different alleles, reducing the probability for correct insertion of the *LoxP* sites) (Figure 3). We therefore hypothesized that success of this approach depends on the combined efficiency of the sgRNA and the likelihood of *LoxP* insertion on both sites to enable two *in cis* HDR events to occur simultaneously. To assess this postulate, we performed a generalized linear regression analysis to model the relationship between Cas9, sgRNA concentration, sgRNA cleavage efficiency, distance between *LoxP* insertions, occurrences of *LoxP* insertions, and success of the 2sgRNA-2ssODN method. The analyses are summarized in Supplementary Table 3. The efficiency of *LoxP* insertions at both 5’ and 3’ sites appears to be the best predictor for the likelihood of success of the 2sgRNA-2ssODN method accounting for over 60% of the total variance.

**Figure 2:**
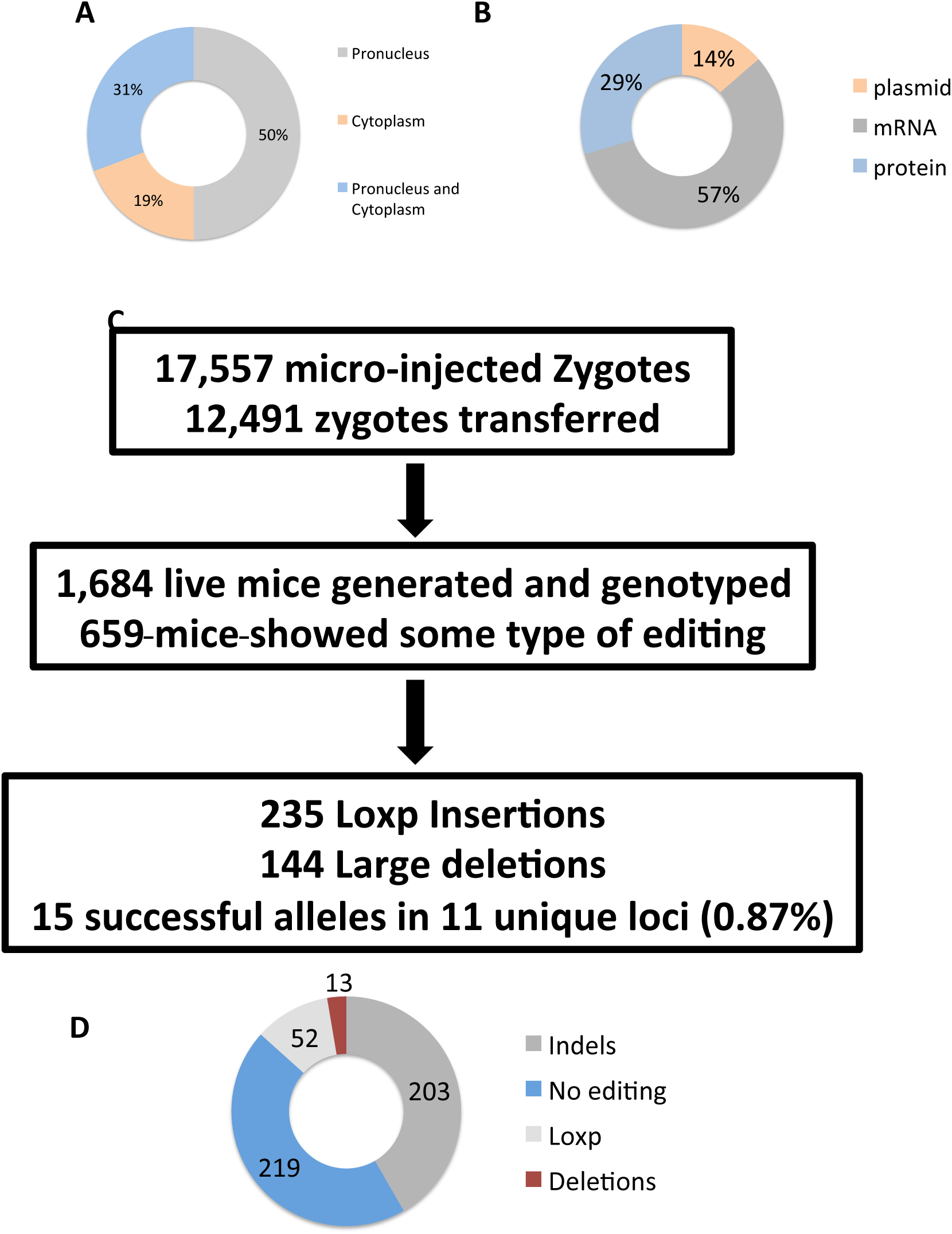
QuanEtaEve assessment of the 2sgRNA92ssODN methods. (A) Doughnut graph indica2ng the methods of zygote injec2ons (pronuclear, cytoplasmic or both) of the CRISPR reagents. Numbers indicate the percentage of the total zygotes microinjected or electroporated. (B) Doughnut graph indica2ng the form of delivery of the CRISPR reagents (mRNA, protein or plasmid) in the zygotes. Numbers indicate percentages. (C) Flow chart indica2ng the number of successful edited alleles and correct *LoxP* inser2ons out of the number of live born pups from microinjected and transferred zygotes. Numbers indicate absolute numbers. (D) Doughnut chart indica2ng the edi2ng types observed amongst the live born pups genotyped on a sub sample from 24 loci. Numbers indicate absolute values.

**Figure 3:**
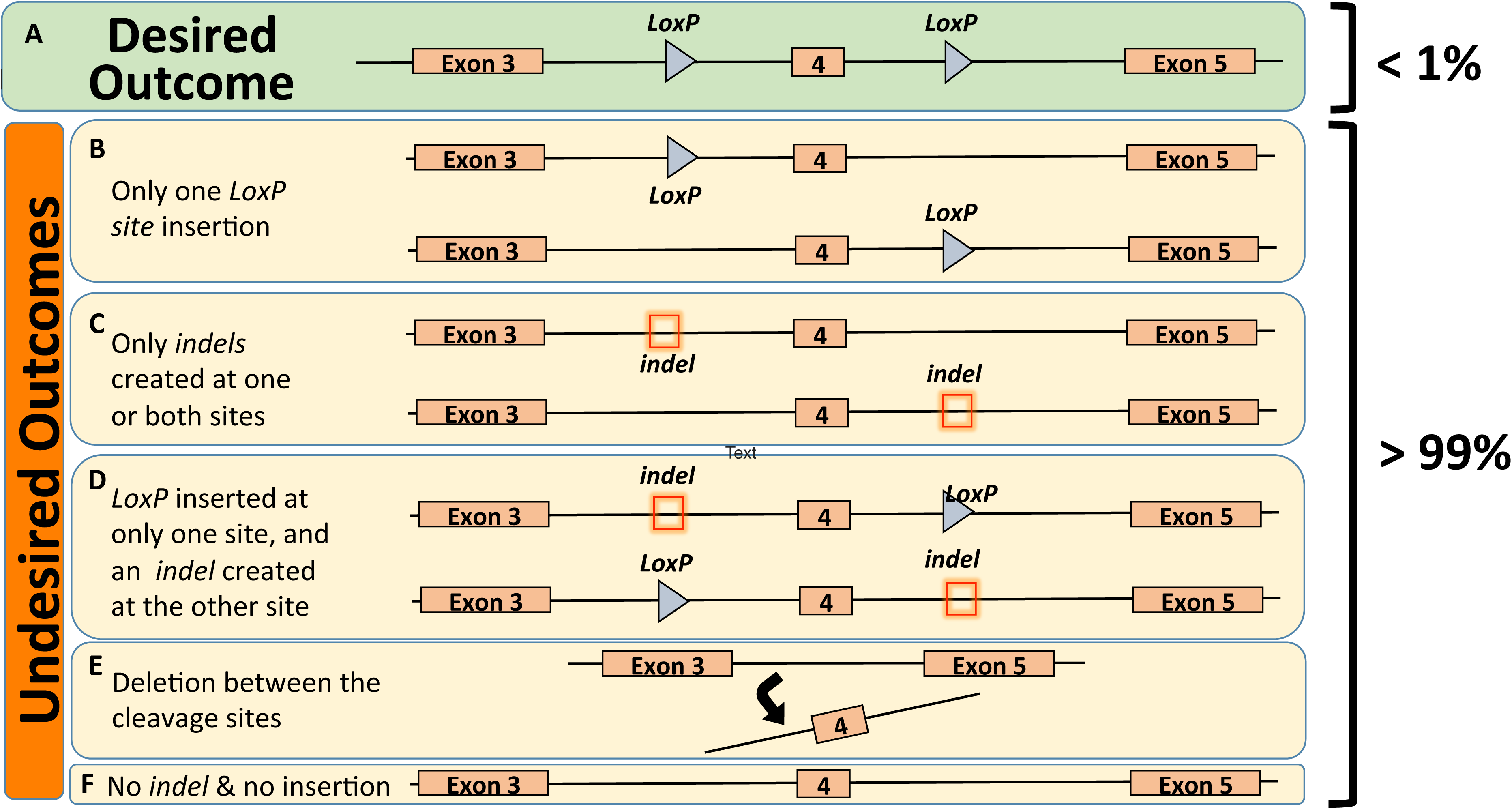
Desired and undesired outcomes of the 2sgRNA-2ssODN CRISPR method of creating conditional knockout alleles. (A). Desired outcome showing a floxed allele and its occurrence is <1%. (B) to (F): various undesired outcomes including only one *LoxP* site insertion (B), only *indels* created at one or both sites (C), combination of *LoxP* insertion and *indels* (D), deletion between the two cleavage sites (E) and no *indel* or no insertion events (F).

However, this predictor was not significant in our linear regression model. Additional predictors such as sgRNA efficiency or efficiency in 5’- or 3’-insertion of *LoxP* explained approximately 35% of the total variance but none of these predictors were significant in our model. The concentration of Cas9 mRNA accounted less than 0.1% of the total variance but was statistically significant (p <0.01) in the generalized linear regression model as a predictor for the success of the 2sgRNA-2ssODN approach. However, success of the 2sgRNA-2ssODN approach was not significantly correlated with an increase of Cas9 mRNA concentration (r^2^ Pearson = 0.27, p=0.08). From our analysis, the sample size of the successful *LoxP* insertions in *cis* was too small to definitively rule out any others predictors (Cohen’s effect size d = 0.4, power 1-beta = 0.41). Together, these results suggest that the presence of two simultaneous HDR events is the best predictor to generating two floxed alleles in *cis*.

Recently, electroporation of zygotes has been developed as an efficient method for generating knockout, point mutations, tagged, or conditional alleles [19-25]. From our consortium, 3 laboratories and programs surveyed the likelihood of success of the method. For 7 loci surveyed, we noted success in inserting a single *LoxP* allele (Supplementary Table 4) from analysis of blastocysts or live mice for 2 out of the 7 loci. In contrast we noted a relatively high frequency of large deletions and *indels* (up to 39% of large deletions) indicating successful editing. However, none of the loci showed two *LoxP* sites inserted in *cis* in the offspring, suggesting that the delivery of CRISPR reagents by electroporation does not make a statistical difference in obtaining a desired outcome from the 2sgRNA-2ssODN floxing approach, although the large numbers of embryos that can be manipulated allows for the recovery of the very small number of those that are correctly targeted.

## Discussion

CRISPR-Cas9 technology has greatly facilitated the generation of mouse lines containing knockout or knockin alleles. However, the generation of conditional alleles remains a challenge using traditional ES cells and CRISPR-Cas9 gene editing technologies. An earlier paper demonstrated 16% efficiency with 2 chimeric sgRNAs and 2 single-stranded oligonucleotides to produce conditional alleles in mice [10].

To evaluate the efficiency of this 2sgRNA-2ssODN method, three laboratories replicated the experiments described in the initial report on *Mecp2* (10) using the same methods to generate the sgRNA and Cas9 and microinjected the mouse zygotes at similar reagent concentrations. Although we observed single *LoxP* site insertions and *indels* at the cleavage sites, the method was unsuccessful in generating two *LoxP* sites in *cis*. A previous report attempting to replicate the findings of Yang et al [10], found an efficiency of floxing *Mecp2* varying from 2% to 8% with the 2sgRNA-2ssODN approach [26]. We surmise the lack of efficiency in targeting *Mecp2* here was due to a low concentration of reagents delivered by microinjection, a notion corroborated by previous work from Horii and colleagues [26]. Of note, it was reported that up to 6% targeting efficiency was achieved using 25 ng/µl of Cas9, 6 ng/µl of sgRNA and 100 ng/µl of ssODN but toxic to embryonic development; these concentrations are 2-fold higher than those described in Yang *et al,* 2013 [10].

### What determines the success of the 2sgRNA-2ssODN method?

For better understand the critical factors predicting the likelihood of success with the 2sgRNA-2ssODN approach, we surveyed 56 unique loci in mice zygote. We noted that the efficiency of simultaneous insertion of the two *LoxP* sites simultaneously was the best predictor of success using this approach. We also noted a low success rate in generating a conditional allele across all centers (< 1%), varying from 0 to 50% (median = 0%) for individual loci. These results are comparable with previous reports demonstrating an important disparity in success rate varying from 0% to 7% of mice harboring two *LoxP* sites insertions in *cis* whether delivered by microinjection [26-29] or by electroporation [26]. We and others also have noted the large number of deletions at the target sites following DNA cleavage [28]. Our results on a larger number of loci suggest the efficiency in generating a successful cKO with the 2sgRNA-2ssODN method is lower than previously described [10]. One hypothesis for this discrepancy in success rate might relate to strain-specific differences. We analyzed this variable and did not find any significant differences among strains, whether the donor strain was a F1 cross, inbred, or outbred mouse line as a donor strain. Another possibility to improve the efficiency of the method is to avoid recombination between the target sites by placing the *LoxP* sites hundreds of kb apart. This was reported previously for a success rate varying from 0% to 18% for 6 loci [30]. We did not find such evidence in our data, although our sample size is too small to formally rule out this hypothesis. A recent report found the successful use of sequential introduction of the *LoxP* sites to improve efficiency and avoid recombination between alleles [26]. Indeed, a 3 to 10 fold improvement in successful targeting was observed, though it should be noted that such an approach requires a more protracted period of time to completion [26]. Additional work has demonstrated over 5 fold improvement in targeting using a long ssODN [22, 29, 31] or double-stranded donor DNA [28].

In conclusion, we find the 2sgRNA-2ssODN method to be inherently biased for *indels* or substitutions at the DSB, deletion between the guide cleavage sites, or *trans* insertion of the *LoxP* sites. Even though the overall success rate is very low — ∼1,200 zygotes were needed to generate 1 correctly targeted animal — it is possible to generate floxed alleles using CRISPR-Cas9 gene editing technology with 2sgRNA-2ssODN. The method, however, requires two inefficient simultaneous HDR events leading to correct insertion of both *LoxP* sites in the *cis* configuration, an outcome we find occurs very infrequently (<1%).

## Acknowledgements

This work was supported by the National Collaborative Research Infrastructure (NCRIS) via the Australian Phenomics Network (APN) (to Gaetan Burgio and Paul Thomas), by an Institutional Development Award (PI: Shelley Smith) P20GM103471 (to CBG, RMQ, DWH, JDE and RR), by NIGMS 1P30GM110768-01 and P30CA036727 (as part of support to University of Nebraska Mouse Genome Engineering and DNA Sequencing Cores), the British Heart Foundation (FS12-57), FS12/57/29717, CH/13/2/30154 and the program grant RG/15/12/31616 (to Kathryn Hentges and Bernard Keavney), the Wellcome Trust grants 107849/Z/15/Z and 105610/Z/14/Z, the Medical Research Council, MR/N029992/1 (to DB and CBL), the National BioResource Project of Ministry of Education, Culture, Sports, Science and Technology/Japan Agency for Medical Research and Development (MEXT/AMED), Japan, the Canadian Institutes of Health Research, MOP#142452 (MCB and LJM). LJM is a member of the Research Centre of the McGill University Health Centre which is supported in part by FQRS. Dr William Thompson was supported by the Indiana Clinical and Translational Sciences Institute, funded in part by grant #UL1 TR001108 from the National Institute of Health (NIH), National Center for Advancing Translational Sciences, Clinical and Translational Sciences Award. D Kent Lloyd is supported from the NIH (UM1OD023221), and work contributed by staff from the UC Davis Mouse Biology Program (MBP) is supported by a grant from the American College of Laboratory Animal Medicine. The work contributed from Xiande Liu, Chad Smith, Eric Jonasch, Xuesong Zhang and Jan Parker-Thornburg is supported from the NIH under the award number P30CA16672. R Sedlacek was supported by LM2015040 (Czech Centre for Phenogenomics), CZ.1.05/1.1.00/02.0109 (BIOCEV), and CZ.1.05/2.1.00/19.0395 by the Ministry of Education, Youth and Sports (MEYS) and by Academy of Sciences of the Czech Republic (RVO 68378050). David Ray was supported from a Wellcome Trust Investigator (107849/Z/15/Z) and the Medical Research Council (MR/P011853/1 and MR/P023576/) grants. Andrew Loudon was supported from a Wellcome Trust Investigator (107849/Z/15/Z), Biotechnology and Biological Sciences Research Council (BB/N015584/1), Medical Research Council (MR/P023576/1). The work contributed from Gloria Lopez-Castejon is supported from the Wellcome Trust (104192/Z/14/Z) and the Royal Society. Pilar Alcaide was supported from the NIH (HL 123658).

---

**Supplementary Table 1.**
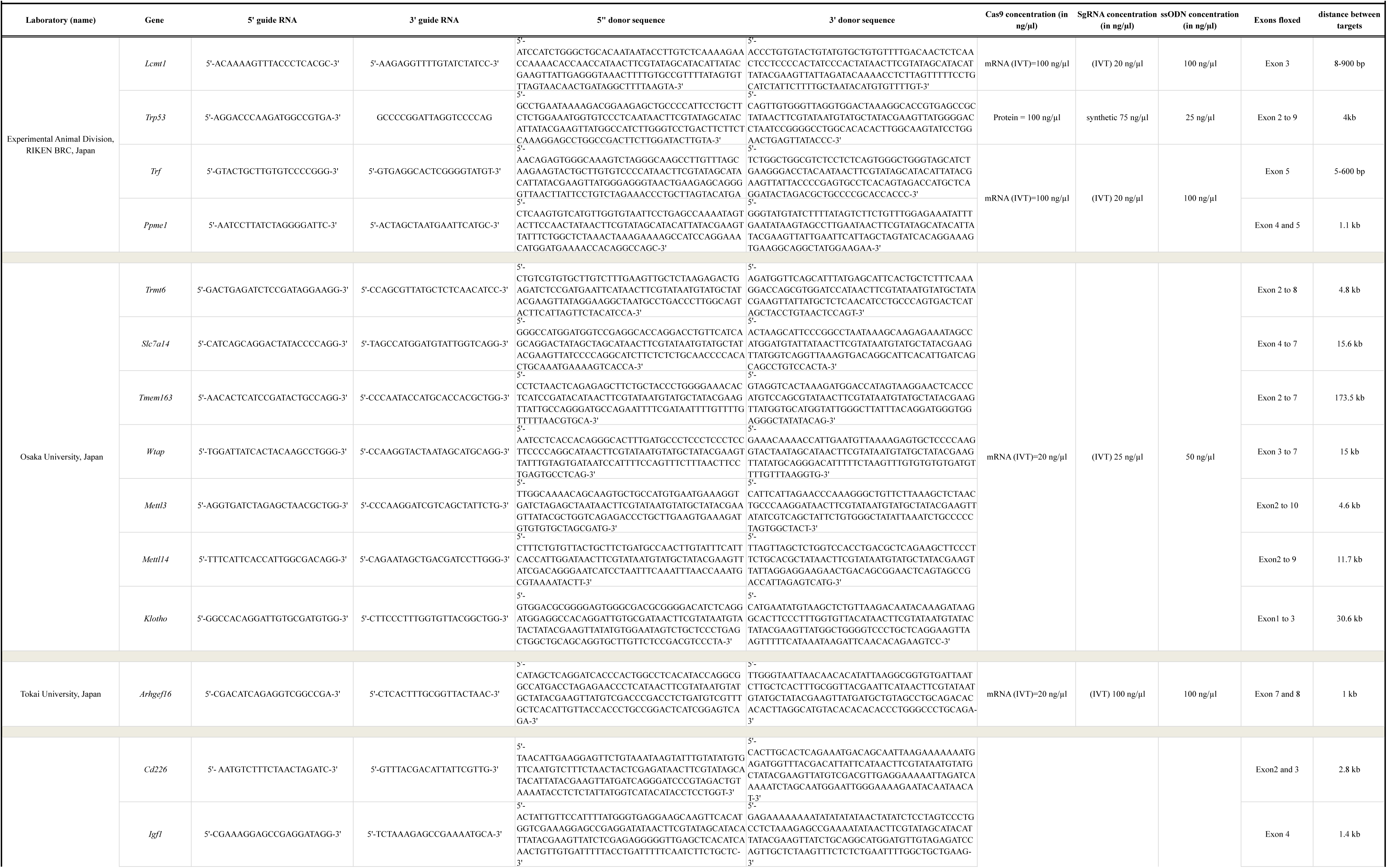
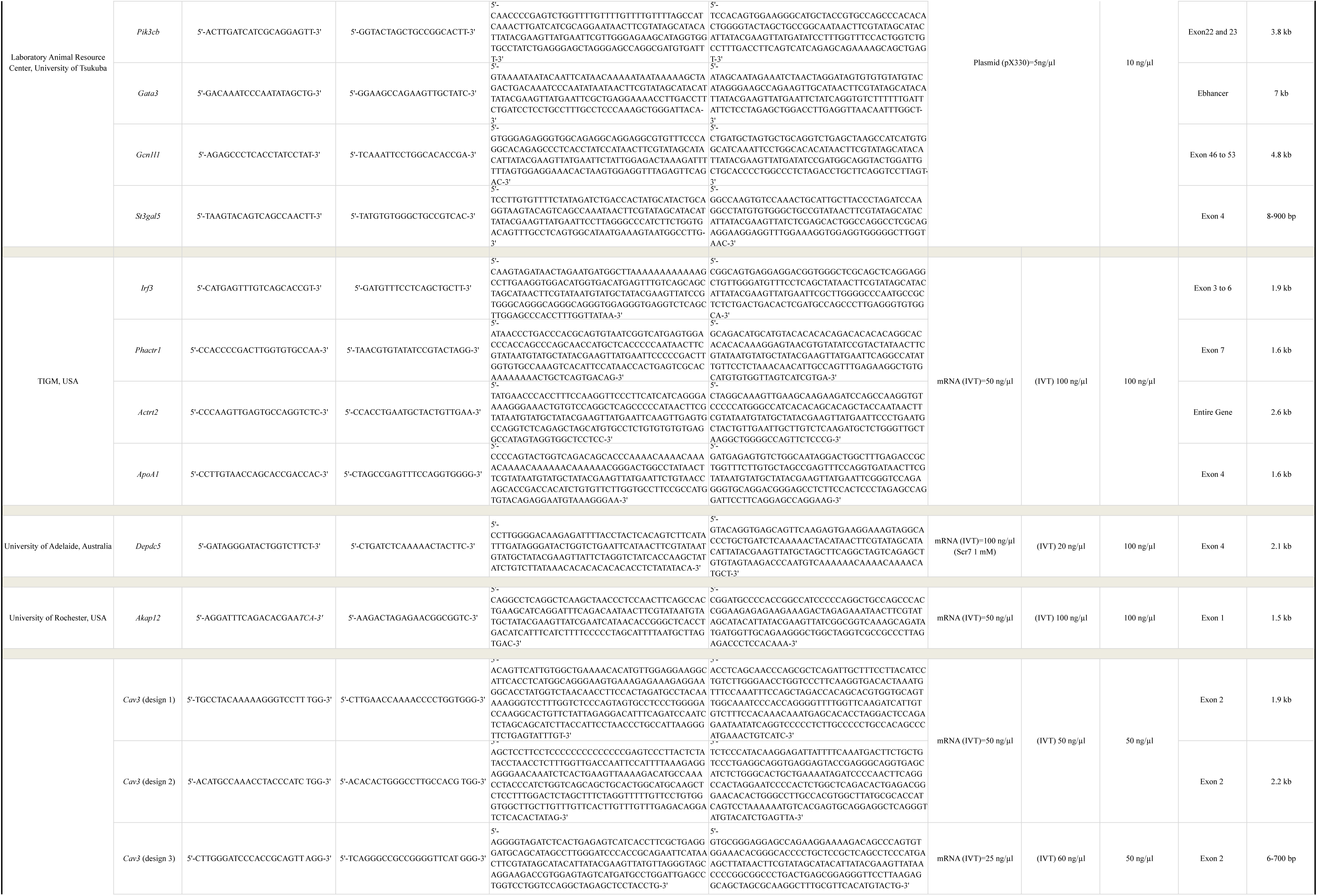
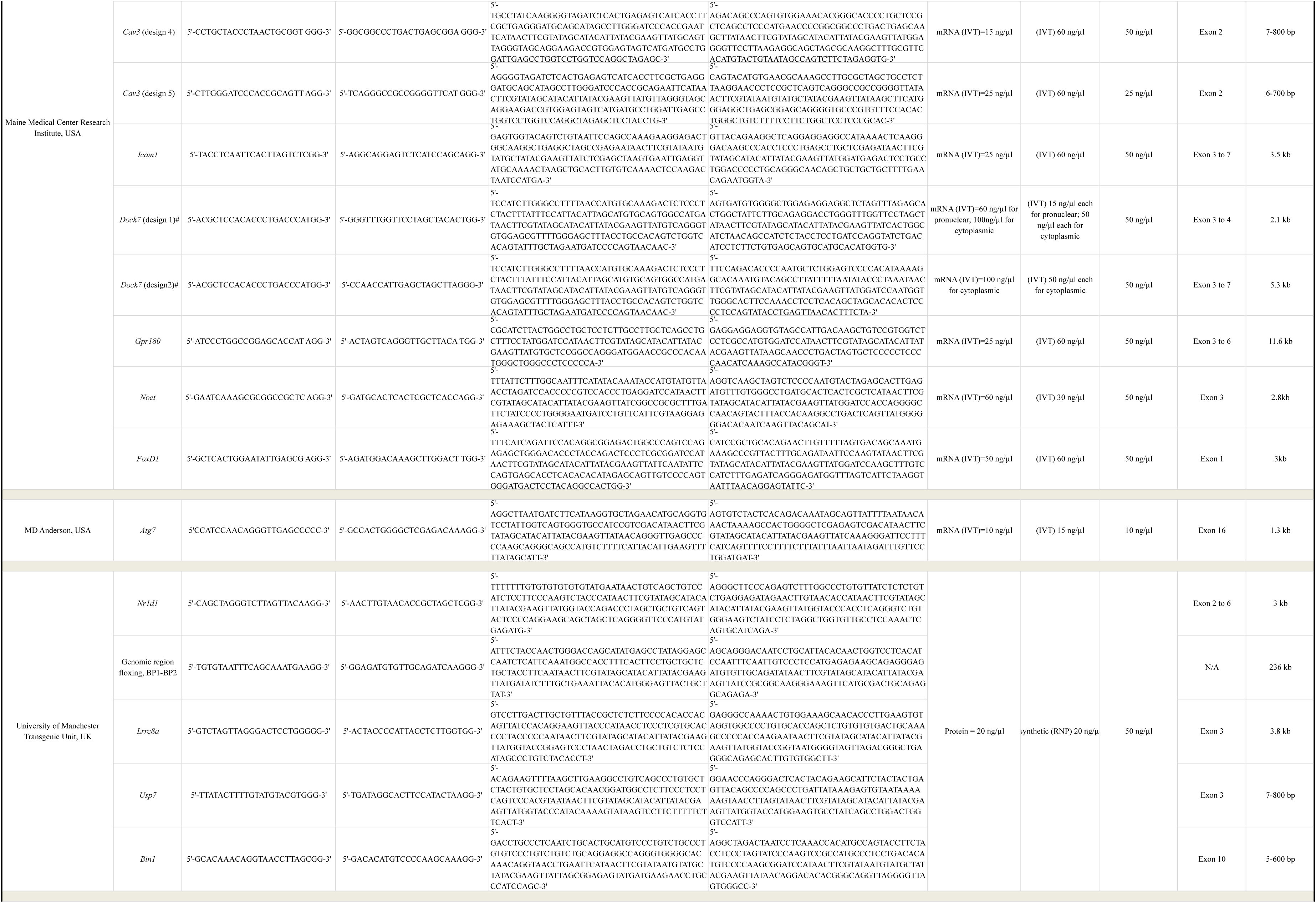
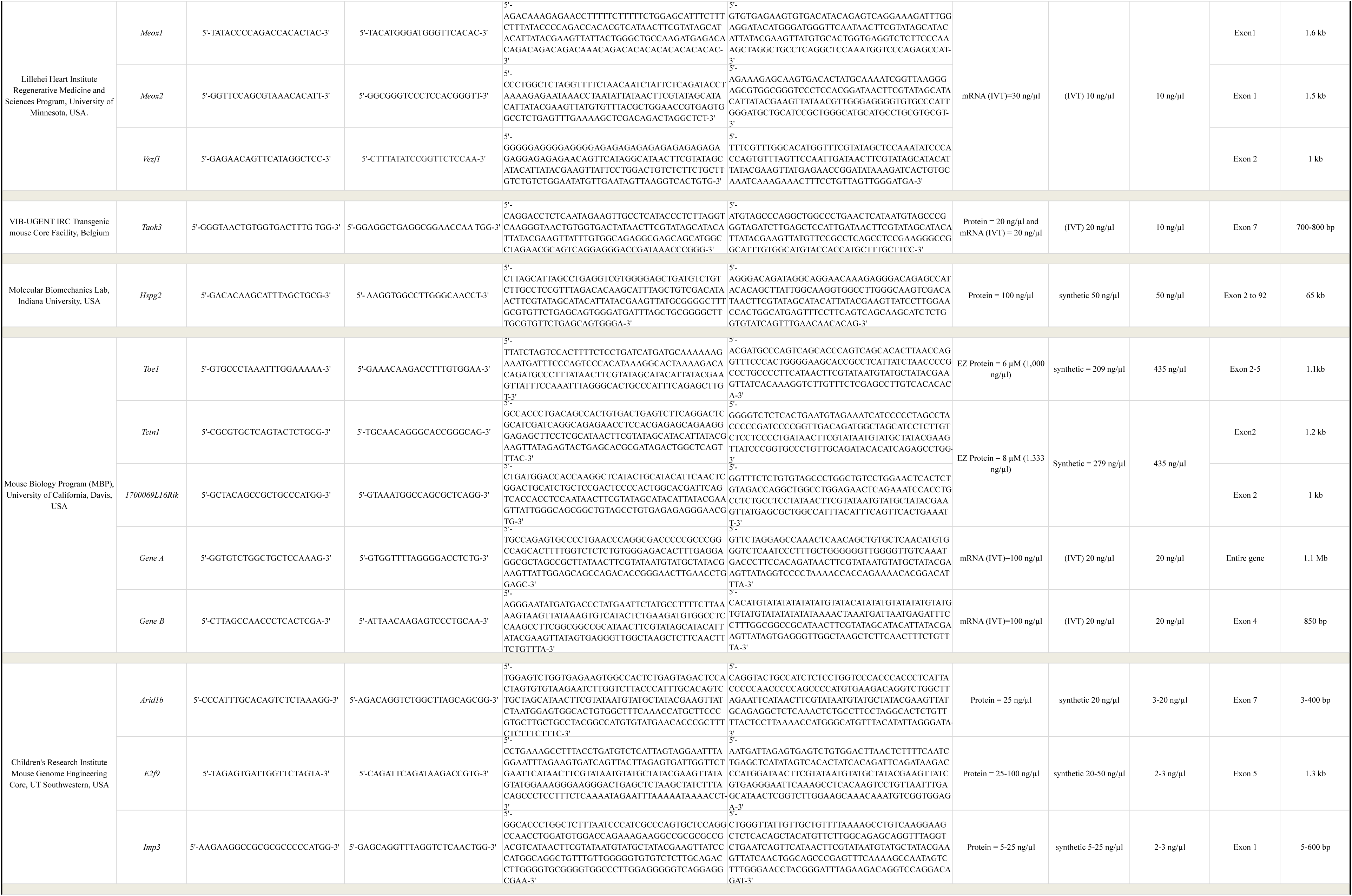
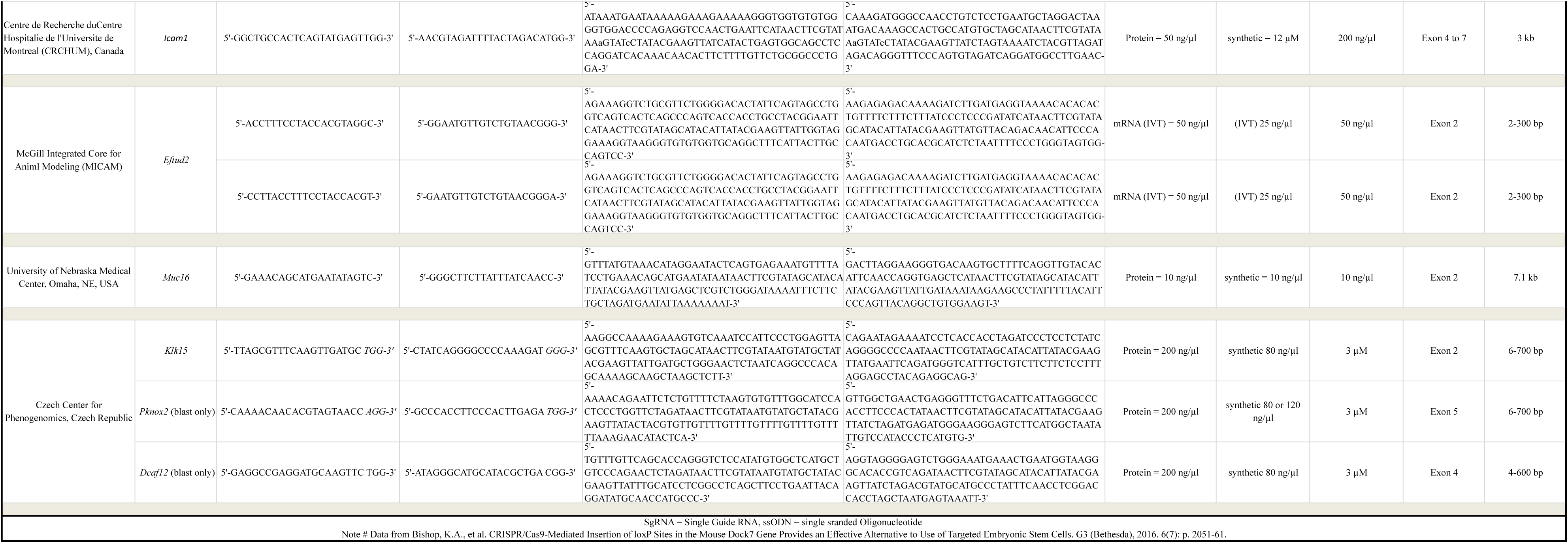
Single Guide RNA, Single Stranded Oligonucleotide DNA sequences, reagents concentrations and targeting genomic regions (in bp) used in this study.

**Supplementary Table 2.**
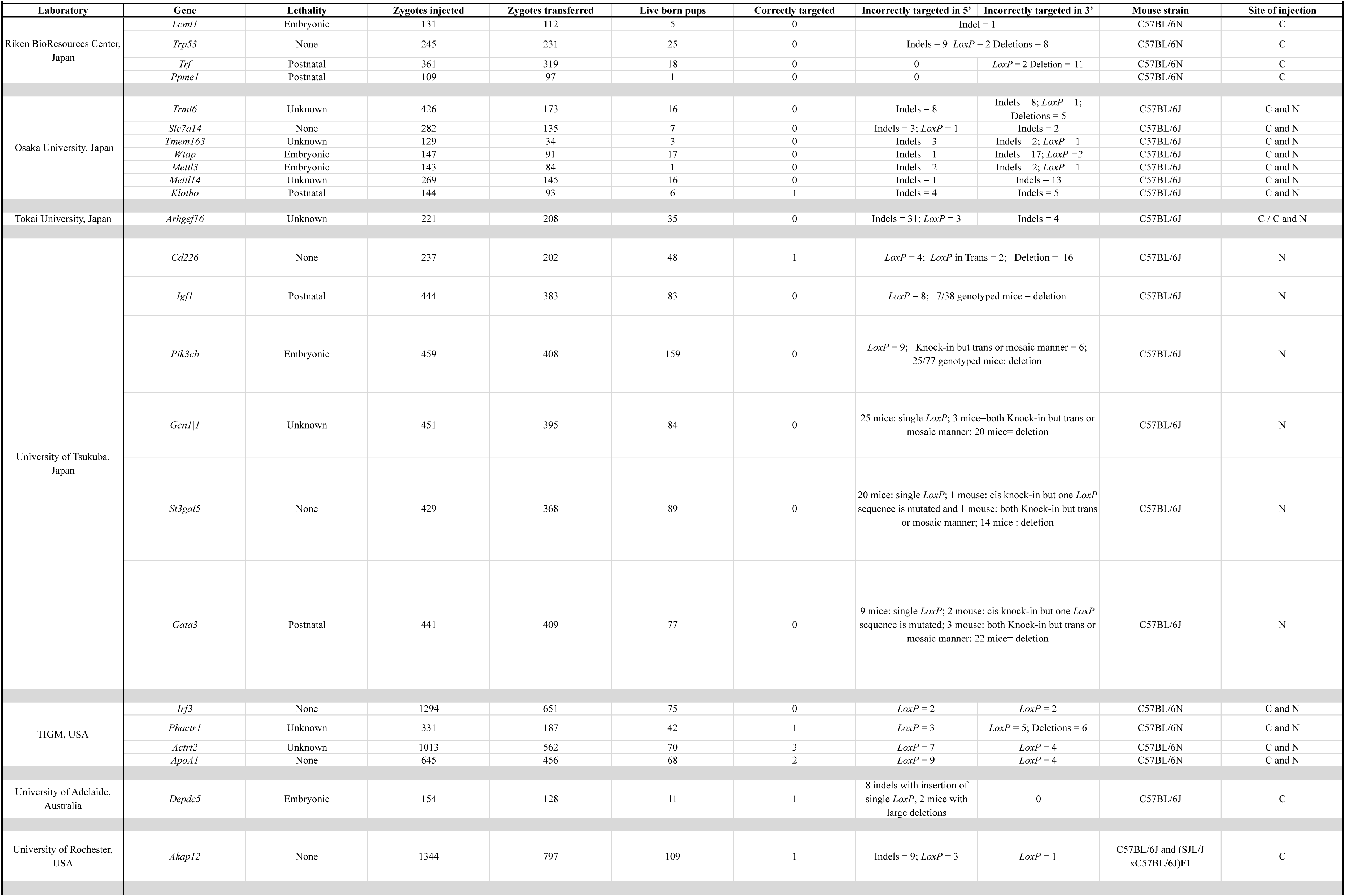
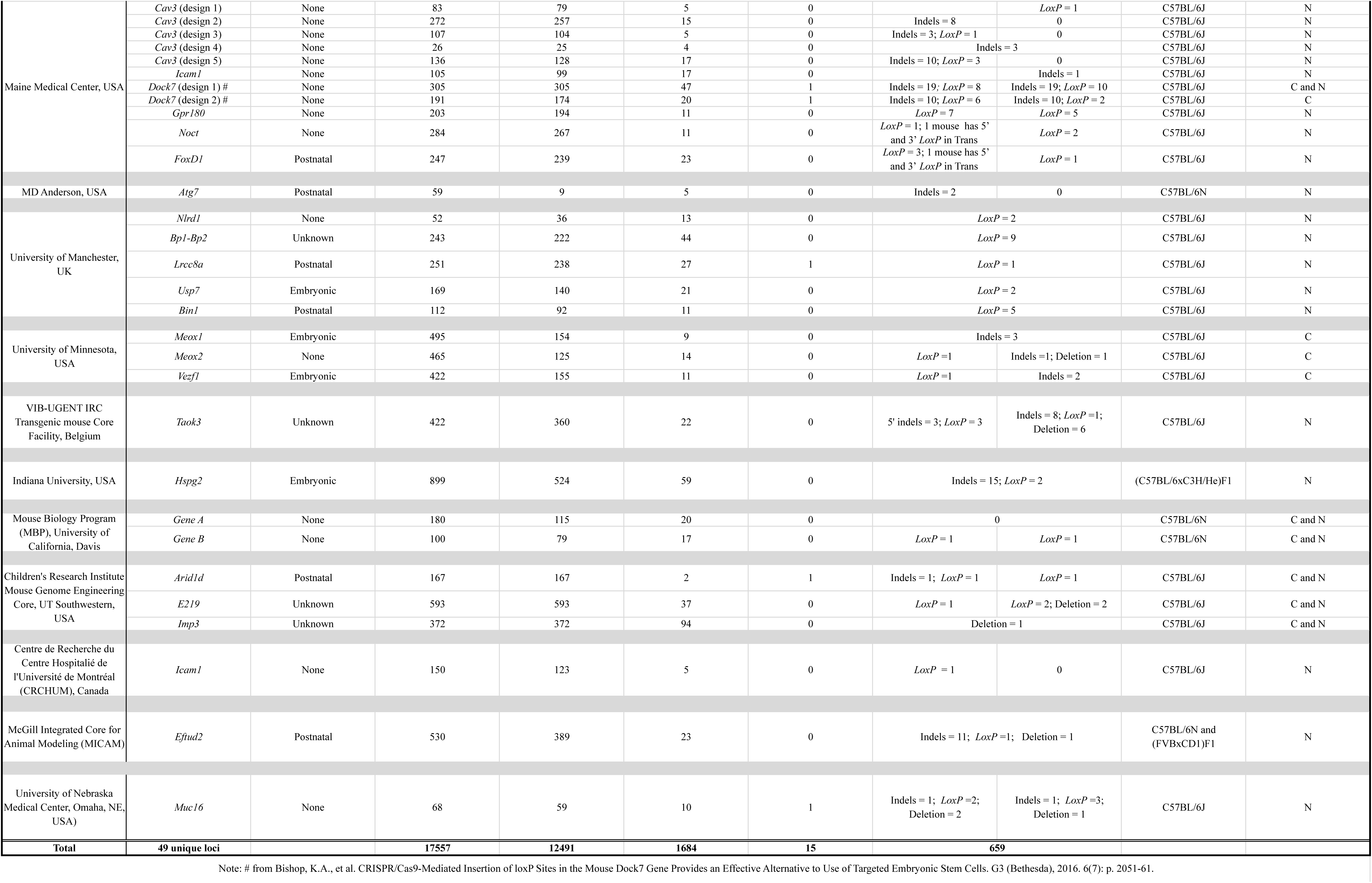
Detailed count of the numbers of zygotes microinjected, transferred, live pups born, correctly targerted and non targeted from 49 unique loci

**Supplementary Table 3.**
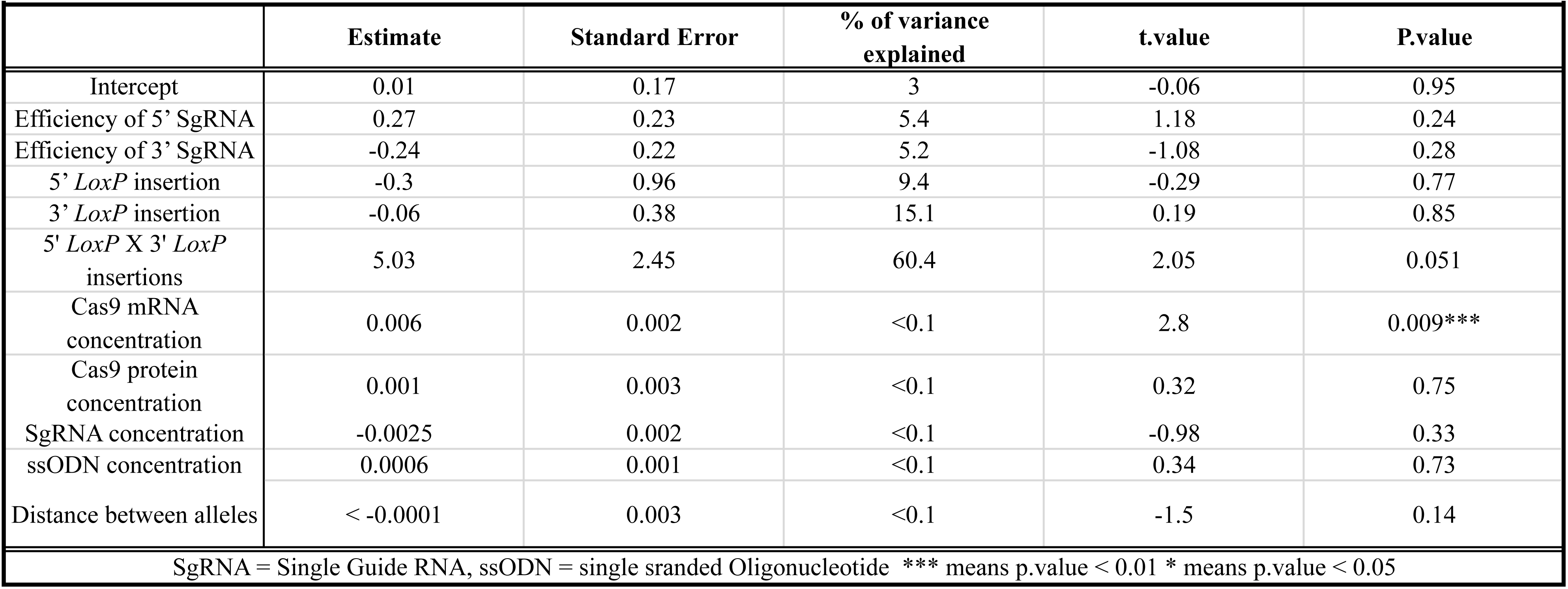
Generalized Regression analysis to identify the factors predicting the success of the 2sgRNA-2ssODN methods

**Supplementary Table 4.**
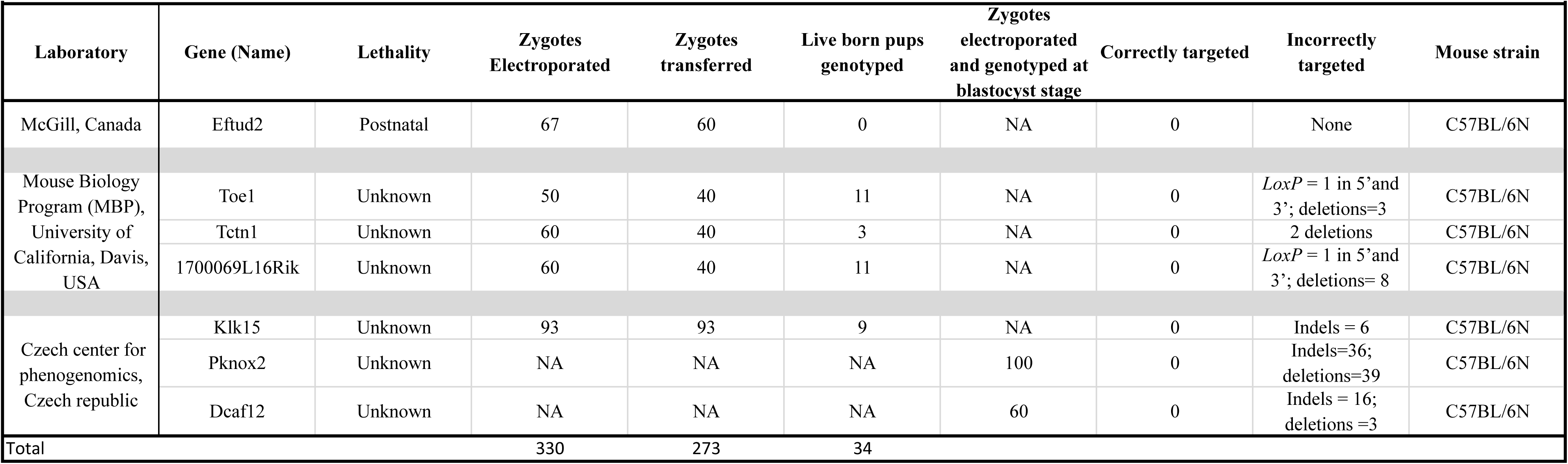
Detailed count of the numbers of zygotes electroporated, live pups born or blastocysts genotyped, correctly targeted and non targeted from 7 unique loci

**Supplementary Table 5.** Overall efficiency of the 2sgRNA-2ssODN method of generating the cKO alleles

|  |  |  |  |  |  |
| --- | --- | --- | --- | --- | --- |
| Delivery method | Number of loci | Zygotes processed | Zygotes transferred (% zygotes processed) | Live born pups analyzed (% zygotes transferred) | Correctly targeted (% live born) |
| Microinjection | 49 | 17,557 | 12,491 | 1,684 | 15 |
| Electroporation | 5 | 330 | 273 | 34 | 0 |
| **Total** | **54** | **17,887** | **12,764 (71.4%)** | **1718 (13.4%)** | **15 (0.87%)** |

